# Joint precursor elution profile inference via regression for peptide detection in data-independent acquisition mass spectra

**DOI:** 10.1101/329805

**Authors:** Alex Hu, Yang Young Lu, Jeff Bilmes, William Stafford Noble

## Abstract

In data independent acquisition (DIA) mass spectrometry, precursor scans are interleaved with wide-window fragmentation scans, resulting in complex fragmentation spectra containing multiple co-eluting peptide species. In this setting, detecting the isotope distribution profiles of intact peptides in the precursor scans can be a critical initial step in accurate peptide detection and quantification. This peak detection step is particularly challenging when the isotope peaks associated with two different peptide species overlap—or *interfere*—with one another. We propose a regression model, called Siren, to detect isotopic peaks in precursor DIA data that can explicitly account for interference. We validate Siren’s peak-calling performance on a variety of data sets by counting how many of the peaks Siren identifies are associated with confidently detected peptides. In particular, we demonstrate that substituting the Siren regression model in place of the existing peak-calling step in DIA-Umpire leads to improved overall rates of peptide detection.

## 1 Introduction

The primary goal of most bottom-up mass-spectrometry-based proteomics is to detect and, ultimately, quantify multiple peptide species on the basis of mass spectra derived from complex biological samples. Each such peptide species is typically observed twice: first in a preliminary (MS1) scan of intact peptides, and subsequently in a secondary (MS2) scan of peptide fragments. Information about the mass and charge of the peptide can be derived from either the MS1 or MS2 data, whereas the specific amino acid sequence of the peptide is typically derived from the MS2 data via database search (reviewed in [13]).

In this work, we propose a method for analysis of MS1 data. Specifically, we focus on the problem of precursor annotation, where a “precursor” is an observed intact peptide ion and the annotation consists of the inferred monoisotopic mass and charge. Such methods are useful in the context of traditional, data-dependent acquisition (DDA) protocols or in the context of increasingly popular data-independent acquisition (DIA) methods. In a DDA setting, precursor annotation analysis helps identify co-eluting precursors that were isolated for an MS2 spectra but do not share the precise m/z of the ions that triggered the sampling of the MS2 spectra, and also may provide better charge state estimates than those based solely on MS2 analysis. MS1 analysis can also help to fix isotope peak errors, in which the mass spectrometer triggers on a peak corresponding to the +1 or +2 isotope, rather than the monoisotope. In a DIA setting, precursor annotation is even more critical, because the isolation window employed in this setting is typically much larger. Hence, in many DIA analyses, the first step toward confidently detecting a given peptide species is to detect its corresponding MS1 precursor.

**Table 1.**
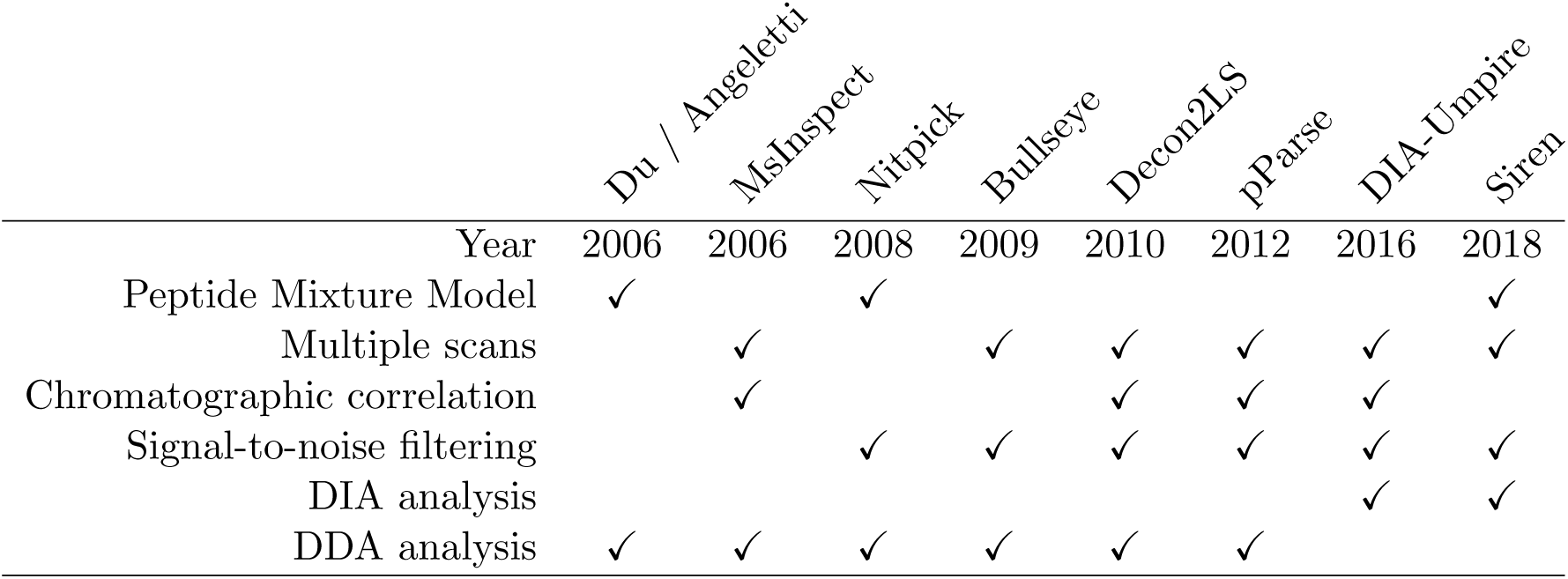
Comparison of precursor annotation methods from MS1 spectra.

### 1.1 Related work

To provide context for our contribution, we first review existing precursor annotation methods. Each of these methods is designed to consider different pieces of relevant information, but no individual method takes all relevant information into account (Table 1). All of the methods rely on averagine models [19] or variations thereof to define models of candidate isotope distributions.

Two early methods use LASSO regression [3, 17] for precursor annotation. The method proposed by Du and Angeletti [3] first identifies candidate theoretical precursors whose two most intense peaks appear in the spectra. The method then uses LASSO regression to narrow down candidates that best explain the observed peaks while trying to avoid false positives. The method Nitpick [17] also uses LASSO to select combinations of candidate precursors, employing a dense, non-centroided representation of peaks and a Bayesian information criterion to eliminate candidates. Both of these regression methods model each observed peak as a sum of ions from multiple peptides at once, allowing the methods to cope with interference. However, neither of the two methods considers information from multiple spectra at once to refine their candidates.

Four other approaches do not employ regression but instead treat each candidate independently. MsInspect [1] chooses candidate precursors by evaluating their peaks over successive scans. This approach allows MsInspect to ensure that its precursor annotations exhibit a characteristic elution profile over time. MsInspect first defines peaks over m/z using wavelet decomposition in each scan and and then filters out candidates that do not persist over multiple scans. The algorithm then evaluates each candidate based on how well the relative intensities of their expected isotope distributions correlate with observed peaks over both m/z and elution time. Bullseye follows roughly the same steps as MsInspect [8]. The first step uses Hardklor [6] to examine MS1 peaks in and around the MS2 isolation window to compute a signal-to-noise ratio and identify which precursor candidates are consistent with non-noise peaks. Bullseye’s second step determines which candidates persist across multiple scans. Decon2LS [9] and pParse [24] also take similar approaches. MsInspect, Decon2LS, and pParse (but not Bullseye) all evaluate candidate precursors based on the correlation of elution profiles of the different isotopic peaks of the candidates.

Software that analyzes DIA data uses and infers precursor information in more diverse ways. DIA-Umpire includes precursor identification as a first step within a larger pipeline to deconvolve MS2 spectra [22]. This method identifies precursor ions by extracting precursor peak chromatograms and identifying them as a precursor’s isotope distribution if exhibit the expected pattern of m/z values and if their intensity profiles correlate well over elution time. DIA-Umpire does not consider observed peaks as mixtures of ions, which makes it vulnerable to interference that disrupts the correlation between isotope peaks over time. Pecan [21] is similar to DDA database search, in the sense that theoretical spectra are scanned against the data, but the score function for each peptide takes into account both MS1 and MS2 data. For the MS1 data, Pecan computes a theoretical isotope distribution for the candidate and scores using the dot product between the theoretical and observed peaks. Because the dot product operation considers only one peptide at a time and is agnostic to the presence of other peptides, this score function will yield a high value for non-existent peptides whose isotope distribution happen to overlap those of existing peptides. Accordingly, Pecan quantifies background signal and revises its interpretation of the dot-product based on the quantification of the background. The method Specter [14] uses regression models to describe mixtures peptides, but it acts at the MS2 level and requires library spectra for every modeled peptide; it does not use MS1 data.

With the exception of the two regression models, all of the subsequent MS1 analysis methods treats each candidate peptide independently. As such, these methods are fundamentally incapable of modeling interference between co-eluting peptides. The regression models, on the other hand, are designed to operate on only a single spectrum at a time.

### 1.2 Contribution

We propose a precursor annotation method called Siren (Sparse Isotope RegressioN) that uses regression to jointly model mixtures of precursors while also taking into account the variation in precursors over time. We provide empirical evidence that Siren’s mixture modeling approach enables proper deconvolution of interference and that the sparse representation allows Siren to efficiently scale to wide DIA isolation windows. Siren also estimates precursor abundances over multiple consecutive scans. Siren is available as open source software implemented in Python <u>(http://bitbucket.org/noblelab/siren)</u>.

## 2 Methods

### 2.1 Linear model of MS1 spectra in DIA data

Siren represents the observed MS1 spectra as a linear combination of theoretical spectra. Siren approximates a single MS1 spectrum as the sum of weighted theoretical precursor isotope distributions (Figure 1a). Repeating this relationship for a sequence of successive scans yields the full model (Figure 1b). Accordingly, the model consists of three matrices.

**Figure 1:**
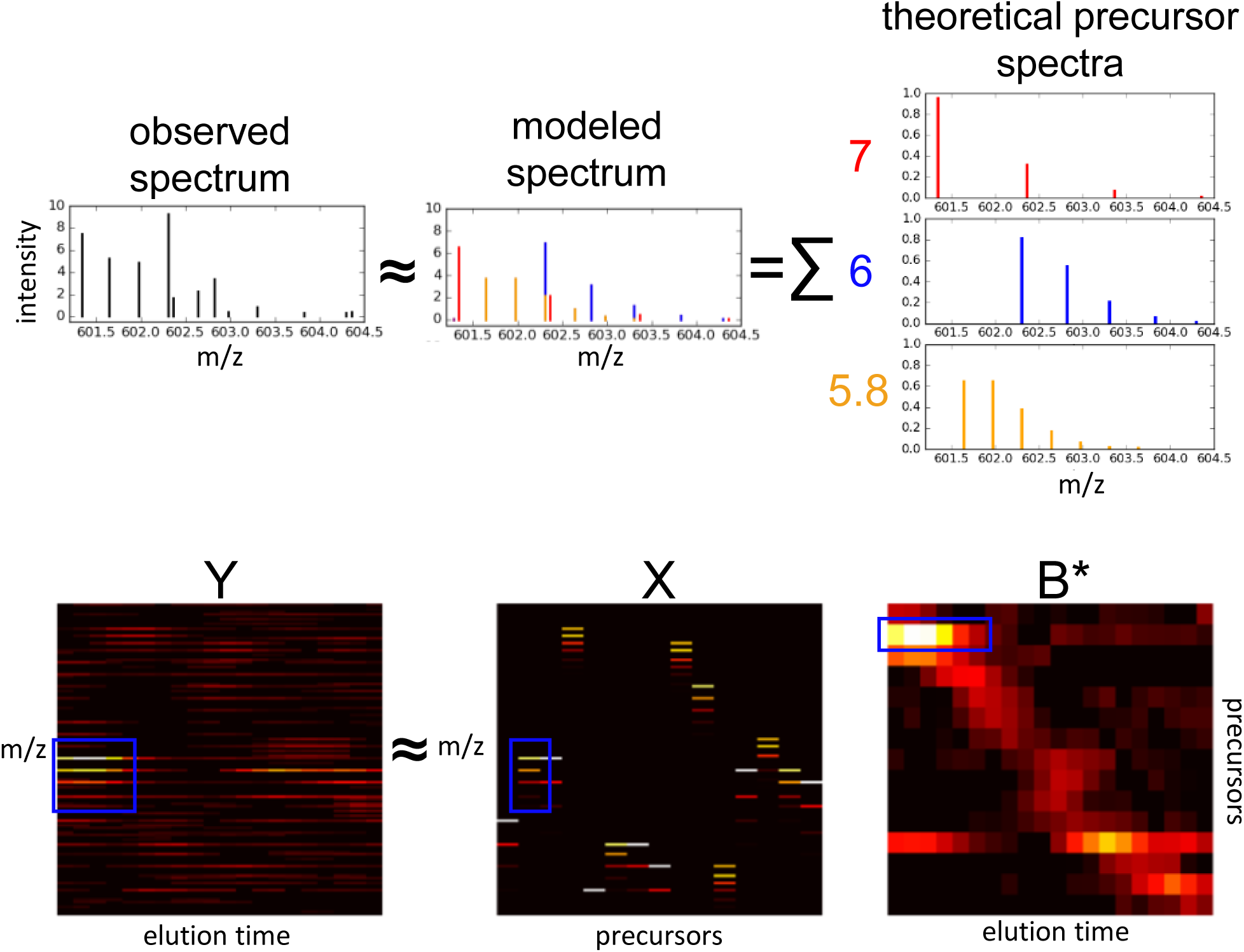
**Siren model.** (A) Each observed MS1 spectrum is approximated by Siren as a linear combination theoretical isotope distributions. (B) Visualization of the Siren approach, using real data. The three heat maps correspond, from left to right, to observed MS1 data, a collection of theoretical isotope distributions, and an inferred abundance matrix. The product of the boxed isotope distribution in *X* the boxed elution profile in *B* yields the boxed pattern visible in matrix *Y*. For clear visualization, the m/z bins in *X* and *Y* are re-discretized, and columns in *X* have been merged to be consistent with *B*.

- *Y* ∈ ℝ ^*MxT*^consists of the observed MS1 data, corresponding to a series of MS1 spectra from a single mass spectrometry run. Each of its *T* columns *Y*_*t*_ is a vector representing a single MS1 spectrum, and each of M rows corresponds to a single m/z bin. Each m/z bin can only be non-zero in a single *Y* column, as each bin is unique to a particular spectrum.
- *X* ∈ ℝ^M*xN*^contains theoretical spectra in the form of isotope distributions for the precursor ions hypothesized to exist in the spectra. Each of *N* columns corresponds to an isotope distribution computed using an averagine model [19] to represent the set of peptide ions that share a particular monoisotopic mass (within a small error) and charge. Note that because the discretized m/z bins are defined separately for each scan, single precursor ion that exists across multiple scans will result in multiple columns, each corresponding to a separate scan and set of m/z bins, whose relationships will be determined in a subsequent step.
- *B* ∈ ℝ;^*NxT*^ contains the inferred abundances of the precursor ions of *X* in *Y*. Each column in *B* represents the abundances of all *N* theoretical precursors in one observed spectrum *Y*_*t*_. *B* is subsequently processed into the matrix *B*\* where redundant rows of *B* that correspond to the same precursor are combined into fewer rows of *B*\*.

Siren assumes that

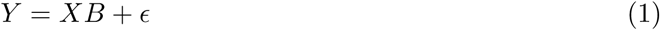

where ∊ represents machine noise, contaminants, and other signals that are not represented by theoretical peaks in *X*. Siren proceeds in three steps: constructing the matrix *Y*, constructing the matrix *X*, and then inferring the matrix *B*. However, Siren exploits the observation that the inference of each column in *B* (i.e., one vector of abundance values across all precursors) can be carried out independently for each column (scan) in *Y*. Thus, the three steps are carried out separately for each scan.

#### 2.1.1 Construction of the observed matrix *Y* and theoretical matrix *X*

Each observed spectrum consists of a set of peaks, with real-valued m/z and intensity values. However, Siren requires that these peaks be arranged in a matrix *Y*. A column *Y*_*t*_ in *Y* corresponds to a discrete scan at discrete elution time *t*, but the rows must be created by some form of discretization of the m/z axis.

Siren does not adopt the conventional strategy, in which the m/z span is divided into equal-width bins, for several reasons. Say that we are working with high-resolution data that exhibits a native resolution of 10 parts-per-million (ppm) on the m/z axis. If we make our bins much larger than 10 ppm, then we end up erroneously placing into the same column peaks that should be assigned distinct m/z values. On the other hand, if we use a fine-grained bin size, then edge effects will introduce arbitrary (and incorrect) distinctions between peaks that truly correspond to the same underlying type of ion (Figure 2). Furthermore, a very fine binning scheme leads to a very large matrix, making calculations very expensive. For example, Nitpick, which was designed for targeted analysis of individual precursors, uses a very fine width of 8 · 10^5^ Th. Using this binning scheme across the commonly used m/z span of 400-1200 Th leads to 10 million bins. Nitpick mitigates off-by-one errors by describing each isotope peak as a Gaussian distribution of many peaks across many m/z bins rather than as a single peak in a single bin, but this smoothing further increases Nitpick’s computational expense.

**Figure 2:**
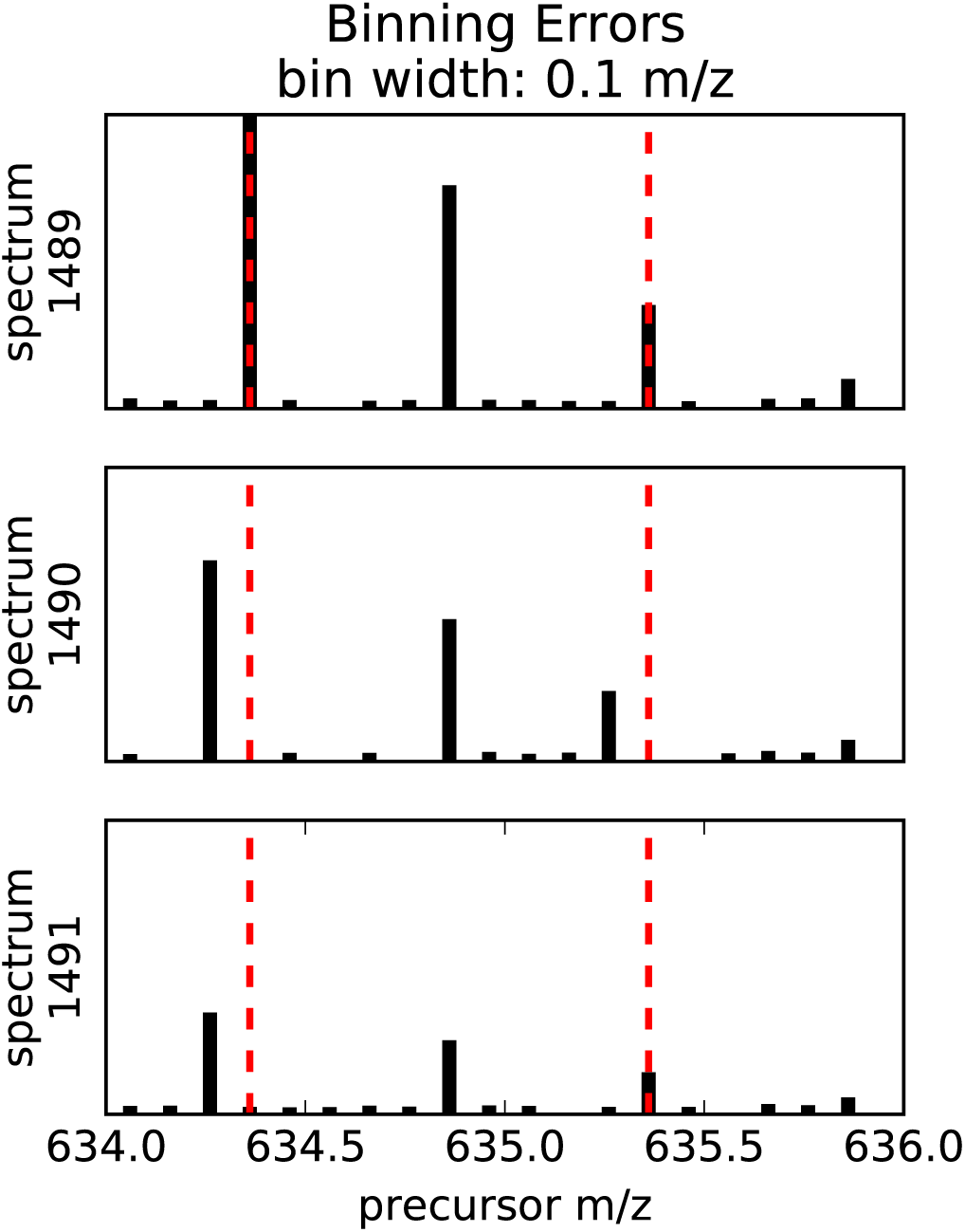
**Binning errors introduced by small m/z bin size.** The figure shows excerpts of a sequence of three MS1 spectra over time, depicting the elution of a charge +2 isotope distribution whose peaks are binned at a 0.1 m/z bin width. The monoisotopic peak and the third isotopic peak each shift one bin to the left from scan 1489 to scan 1490.

Rather than arbitrarily binning the m/z axis, Siren takes an approach similar to Du and Angeletti [3] by creating entries in *Y* that correspond to observed monoisotopic peaks and their corresponding isotope peaks. These entries sparsely describe the important portions of the m/z range, while ignoring m/z values unoccupied by informative peaks. The specific procedure to create a vector *Y*_*t*_ for a single observed scan *T* is as follows:

1. **Noise reduction** Peaks that occur in a single scan, with no corresponding peaks in neighboring scans, likely represent noise. Accordingly, for a scan at time *T* and a peak with m/z value m, we consider the scans at times *T* + 1 and *T* — 1. The current peak is eliminated if neither of the adjacent scans contains a peak with m/z value in the range [m — τ, m + τ], where τ is an m/z tolerance value reflecting the precision of the data.
2. **Observed peak clustering** Peaks are clustered together if they fall within a noise m/z tolerance, with the assumption that small variations in m/z value represent imprecise measurements of the same ion species that should be represented in the same row in *Y*_*t*_. The peaks are represented as a graph, in which vertices are peaks and an edge connects pairs of peaks within a small m/z tolerance of each other. Peaks are then clustered, such that each cluster corresponds to a connected component within the graph. Each cluster a has an associated range of m/z values, *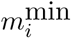* to 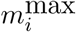.
3. **Monoisotope identification** For each peak cluster *C*_i_, we look for a sibling peak cluster that is separated from *C*_i_, by an m/z difference associated with a pair of isotopic peaks. For a pair of clusters *C*_i_, and *C*_*j*_, a sibling relationship is identified if and only if the m/z difference between the clusters is consistent with the distance between the first two isotopic peaks of a precursor of charge +1, +2, +3 or +4. Formally, the latter condition corresponds to

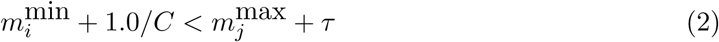

and

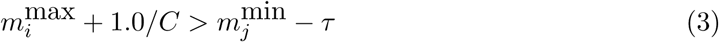

for charge values of C ∈ {1, 2, 3, 4}. Each time a sibling cluster is identified, the initial cluster is defined as the tuple (*m*_*i*_,*C*), and (*m*_*i*_,*C*) is added to *P*_*t*_ to denote the hypothesis that a precursor of mass mi and charge *C* is in spectrum *t*. This step ensures that only monoisotopic precursors whose first two isotope peaks exist in the data are included in the model. Note that, in this procedure, a single cluster can occasionally be marked with multiple charge states. Further charge-state discrimination is performed by regression described below.
4. **Theoretical peak generation** In principle, the m/z and relative intensities of peaks in an isotope distribution can be accurately predicted from the charge of the ion and its elemental composition [18]. However, in our setting, the elemental compositions of each ion in the sample is not known a priori. Accordingly, Siren employs the “averagine” model to approximate a theoretical isotope distribution for each observed monoisotopic peak with m/z *m*_*i*_ = 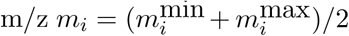 and charge *C*_*i*_ [19]. The calculation retains the +1 through +6 isotope peaks, each with an m/z value and a relative intensity.
5. **Construction of** *Y*_*t*_. The vector *Y*_*t*_ is constructed with one entry corresponding to each observed monoisotopic peak, plus additional entries for the +1 through +6 peaks in each marked charge state. Intensity values for the monoisotopic peaks are directly observed. For the remaining isotope peaks, if an observed peak exists in the data within an m/z tolerance of τ, then the observed intensity is used; otherwise, the intensity is set to zero.
6. **Construction** of *X*. The matrix *X* of theoretical isotope distributions is constructed such that rows in *X* correspond to entries in *Y*. The averagine isotope distributions are placed into *X*, scaled so that the L2 norm of each column in *X* is equal to 1. If the theoretical isotope peaks across *X* do not exist within τ m/z of an observed peak, then they are clustered with each other in the same way as described in Step 2. Each of these clusters forms a row in *Y* and *X*.

#### 2.1.2 LASSO regression to identify precursors

Finally, given matrices *Y* and *X*, Siren assumes the existence of a “true” abundance matrix *B\** and estimates it by solving the following optimization problem:

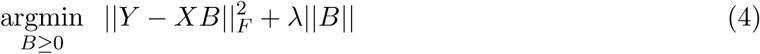

This is known as LASSO regression with a non-negative constraint [20]. The first term is the ordinary least-squares penalty that penalizes differences between the observed data *Y* and the model of the observed data *XB*. The optimization problem balances the first penalty’s goal to accurately model the data against the second term that penalizes the absolute value of *B* to control overfitting.

The parameter λ controls the relative weights of the two penalties. The solution *B* to the optimization problem for a given λ is an estimate of *B*^*^. Increasing λ decreases the inferred abundances of the peptides and infers the presence of fewer peptides overall, thereby decreasing the likehood of false positive inferences at the expense of underestimating abundances and increasing false negative inferences.

Siren uses the algorithm Least Angle Regression (LARS) modified to compute LASSO solutions [4] for every value of λ (the L1 regularization path), implemented in Scikit-learn [15], to compute *B* for every non-negative value of λ.

#### 2.1.3 Extracting continuous elution profiles from B

Further processing must be done on the inferred *B* to form continuous elution profiles because of the way *X* and *Y* are constructed, separate rows of *B* may actually correspond to the same precursor ion hypothesized to exist separately in consecutive scans. To form continuous elution profiles for each precursor, non-zero values in *B* that correspond to approximately the same mass, the same charge, and are adjacent in time are stitched together to form continuous elution profiles. These continuous elution profiles form the matrix *B\** with *T* columns and *N\** < *N* rows, where *N\** is determined after the *B\** is constructed by the algorithm below. The decoy columns of *X* and their rows in B^*^ are constructed separately but using the same algorithm.

1. The vector *B* is designated as the first the column of *B*\*.
2. Subsequent vectors *b*_*t*_ are appended one by one to *B*\*. To add *b*_*t*_ to *B\** that already contains {*b*,…, *b*_*t*-1_}, each value *b*_*n*_,_*t*_ in *b*_*t*_ is appended to a row of *B*\*:

a. If the monoisotopic m/z of *b*_*n*_,_*t*_ overlaps with that of a precursor of the same charge already in *B\** that is non-zero at time *t—* 1, then *b*_*n*_,_*t*_ is appended to that precursor’s row in *B*.
b. If the monoisotopic m/z of *b*_n_,_*t*_ overlaps with more than one non-zero precursor of the same charge already in *B*, then the rows for the overlapping precursors are summed together and merged into a single row in *B*, and *b*_*n*_,_*t*_ is appended to that row.
c. If the monoisotopic m/z of *b*_*n*_,_*t*_ does not overlap any precursor of the same charge with a non-zero value at time *t* — 1, then a new row of zeros is created in *B* and assigned to *p*_*t*_,_*n*_, and *b*_*t*_,_*n*_ is appended to that row.
3. After the entirety of *b*_*t*_ has been added to *B*\*, a zero is appended to all rows that did not get assigned a new precursor from scan *t*.

#### 2.1.4 Identification of precursor elution profiles in *B*\*

Each row of constructed matrix *B\** of inferred abundances may contain elution profiles from multiple peptide ions that share the same mass and charge. To identify individual elution profiles from a row of *B*\*, we use the following procedure. For each precursor (row) in *B*\*, we apply Savitzky-Golay smoothing (using a third degree polynomial over five points at a time) [16]. Local maxima are then found in the smoothed row as points that are greater than both of its adjacent values. We only consider maxima whose adjacent values are both greater than 0. Each local maximum is considered to be an elution peak for a single precursor, and the elution profile of that precursor is bounded by the adjacent local minima.

#### 2.1.5 Estimating the false-discovery proportion of inferred precursors as a function of λ

A proportion of the inferred elution profiles will be incorrect because of noise, non-peptide contaminants, and inaccuracies inherent to the averagine model of peptide isotope distributions. The proportion varies by the value of *λ*. To estimate this false-discovery proportion (FDP), columns containing decoy isotope distributions are included in *X* to facilitate the estimation. For each real (target) theoretical isotope distribution in *X*, a decoy distribution is generated by reversing the order of the intensities of the isotope peaks in the target distribution while preserving the m/z of each peak, and each decoy distribution is added as a column in *X*. Subsequently, for a given value of λ, we can then estimate the FDP as the proportion of inferred elution profiles associated with decoy isotope distributions in *B*.

### 2.2 Data sets

The first data set is derived from HEK-293 lysates using an Orbitrap Fusion mass spectrometer over a 135-minute gradient [23]. Data was downloaded as mzXML files from ftp://ftp.pride.ebi.ac.uk/pride/data/archive/2016/06/PXD003179/. The two runs used are referred to as Tsou2016A with 2205 MS1 spectra (B_D140314_SGSDSsample1_R01_MHRM_T0.raw), and Tsou2016B with 2219 MS1 spectra (B_D140314_SGSDSsample1_R.02_MHRM_T0.raw). The isolation widths vary from 24 to 222 m/z across the range 400-1250 m/z.

The second data set, Navarro2015, is derived from mixtures of lysates from three sources, human, yeast, *E. coli,* and human, mixed in a ratio of 13:6:1, respectively [12]. This data set was used as a benchmark to evaluate peptide identification and quantification algorithms. These experiments were run on a TripleTOF 5600 mass spectrometer with a 120-minute gradient. Data were downloaded as wiff files (HYE110_TTOF6600_32fix_lgillet_I160308_001.wiff) from ftp://ftp.pride.ebi.ac.uk/pride/data/archive/2016/09/PXD002952. This data set contains 2197 MS1 spectra with fixed m/z isolation windows of 25 m/z spanning from 400 to 1200 m/z.

The third data set, Bruderer2017, is derived from human spinal cord tissue and was generated on a Q-Exactive mass spectrometer over a 120-minute gradient [2], downloaded as a raw file from PeptideAtlas (PASS00782). This data set contains 2228 MS1 spectra, and employs precursor isolation widths that vary from 24 to 222 m/z across the range 400-1220 m/z.

All files were converted to .ms1/.ms2 (for Siren) and .mzXML (for DIA-Umpire) formats using msconvert [10].

For all three data sets, downstream analyses were carried out using a database of human tryptic peptides derived from Uniprot (11/02/2014).

### 2.3 DIA-Umpire analysis

We use DIA-Umpire’s deconvolution pipeline to identify precursor elution profiles and deconvolve the MS2 spectra into one pseudospectrum for each elution profile. DIA-Umpire categorizes pseudospectra into three sets: Q1 if its first three precursor isotope peaks appear in the MS1 spectra, Q2 if the first two appear, and Q3 if only its precursor peak appears in the MS2 spectra. We only report analyses for Q1 precursors, as the addition of Q2 and Q3 precursors in the Tsou2016 dataset result in fewer confident peptide identifications. Note that, for the raw MS2 analysis (Section 3.2), we ignore the deconvolved MS2 information within the pseudospectra and extract only the precursor masses, charges, and peak elution times. With this information, we extract corresponding raw MS2 spectra for input to Tide.

### 2.4 Tide analysis

We employ the Tide search engine with exact p-values [7] to search raw (Section 3.2) or deconvolved (Section 3.3) MS2 spectra. The precursor isolation widths were set to 0.01 m/z for the Tsou2016 data sets, 0.01 m/z for the Navarro2017 data set, and 0.01 m/z for the Bruderer2017 data set. Shuffled decoys were created by Tide, and searches were carried out in concatenated mode to allow target-decoy. Peptide-level FDR was estimated using the “weed-out then estimate” procedure [5] using in-house scripts.

### 2.5 Software implementation

The Siren software is implemented in Python and is publicly available at http://noble.gs.washington.edu/proj/siren with an Apache license. The software depends on the packages numpy, scikit-learn, and scipy. Siren takes as input precursor spectra in the form of .ms1 files [11] and outputs identified elution peaks in a text file.

## 3 Results

We tested Siren’s performance in three ways: how accurately its regression model can describe a real MS1 data set, how many peptides associated with its precursor annotations can be identified from raw MS2 spectra via database search, and how many peptides can be identified when Siren is incorporated into DIA-Umpire. Siren has a single tunable parameter λ that effectively controls the number of peptides detected, so we also investigated the effect of this parameter on performance.

### 3.1 Ability of theoretical isotope distributions to model the observed data

For Siren’s regression modeling approach to make sense, it must be the case that a real MS1 data set *Y* can be accurately decomposed into two matrices *X* and *B*. To test this property, we applied Siren to a previously described DIA data set (Tsou2016A) derived from the human cell line HEK-293 [23]. As a control, we created a shuffled version of the Tsou2016A data set, in which the intensities within each scan are shuffled uniformly at random among all the peaks in that scan. For each data set, we use Siren model to infer an abundance matrix *B*. Note that, for this analysis, we set λ = 0 to maximize the model’s accuracy in modeling peak intensities; non-zero values of λ result in the underestimation of the values in *B* and the modeled peak intensities in *XB*.

This analysis shows that Siren’s model is indeed capable of capturing most of the empirical structure of the Tsou2016A data set. We quantify performance using the average R^2^, defined as

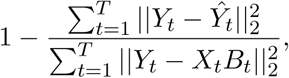

Where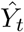 is a matrix filled with the mean value in *Y*_*t*_. of the same dimension as *Y*_t_.We observe that the learned models produced an average *R*^2^ of 0.977 over all MS1 spectra, meaning that 97.7% of the variance in peak intensities was accounted for by the Siren models on average. Furthermore, as expected, Siren’s average *R*^2^ on the shuffled control data set decreases, from 0.982 to 0.602. These results imply that real data can be accurately described as linear combinations of precursor isotope distributions, and that the necessary structure is lost under randomization of peak intensities.

Next, as a control, we created a shuffled version of the Tsou2016A data set, in which the intensities within each scan are shuffled uniformly at random among all the peaks in that scan. As expected, because this shuffled data does not exhibit the expected structure, Siren’s average *R*^2^ decreases, from 0.982 to 0.602. This result implies that the data can be accurately described as linear combinations of precursor isotope distributions.

### 3.2 Comparison of Siren and DIA-Umpire using raw MS2 spectra

Next, to test the quality of the peak calls produced by Siren, we aimed to compare its performance to that of Bullseye, Nitpick, and the peak-calling component of DIA-Umpire, again using the Tsou2016A data set. Unfortunately, we found that Nitpick, which was designed for targeted analysis of individual precursors, cannot scale to analysis of a full MS1 run. Also, Bullseye, which is no longer actively maintained, yielded very few validated identifications. Therefore, our analysis focused on a comparison to DIA-Umpire, which is a recently developed and actively maintained tool.

We used MS2 data to evaluate the quality of peaks called by each method, by counting the number of candidate peaks that successfully lead to a detected peptide in a downstream database search. For this analysis, the database search is conducted using raw MS2 spectra from the DIA data set. This approach is clearly problematic, because the large precursor window used to generate each MS2 spectrum will lead to many co-eluting peptides and hence low overall statistical power in the database search step. However, we reasoned that the “handicap” induced by the use of raw MS2 spectra would be equal for the two methods. In Section 3.3, we report results of a similar analysis conducted using deconvolved MS2 spectra.

Overall, DIA-Umpire predicts a larger number of peaks than Siren (Figure 4)A. For this data set, DIA-Umpire yields > 160, 000 pseudospectra, whereas even at its most relaxed setting (λ = 0), Siren only assigns non-zero abundances to 69% of the hypothesized precursors, leading to ^130,000 elution peaks. When we increase λ, the number of peaks produced by Siren drops substantially. Surprisingly, irrespective of the λ threshold, the overlap between peaks assigned by Siren and peaks assigned by DIA-Umpire is quite low.

**Figure 3:**
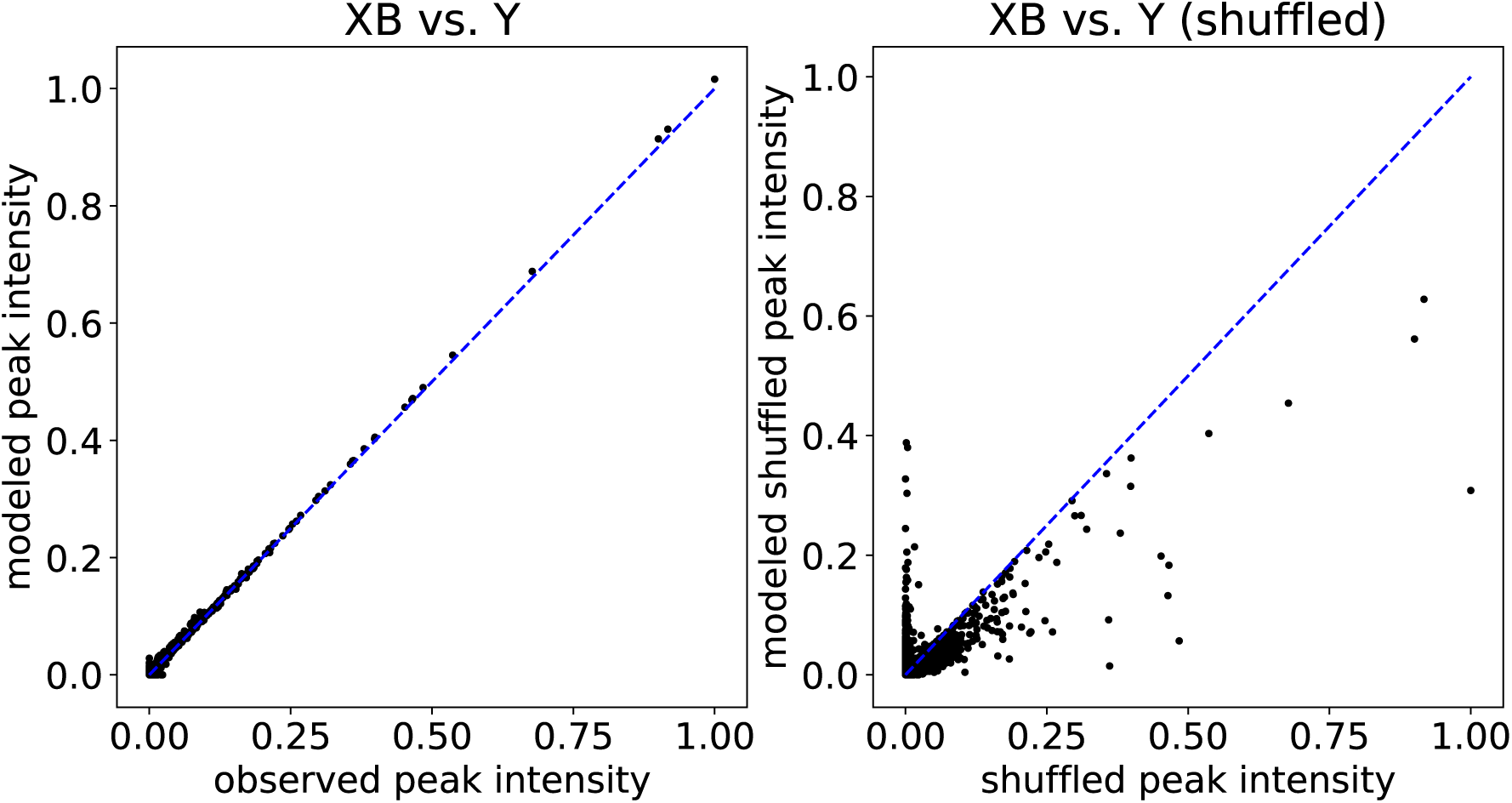
**Observed versus modeled peak intensities from Siren.** Each point in the plot corresponds to an MS1 peak, with its true intensity (x-axis) versus its intensity as modeled by Siren (y-axis). The left plot shows results for peaks from 18 spectra, and the right plot is for peaks shuffled from those 18 spectra.

**Figure 4:**
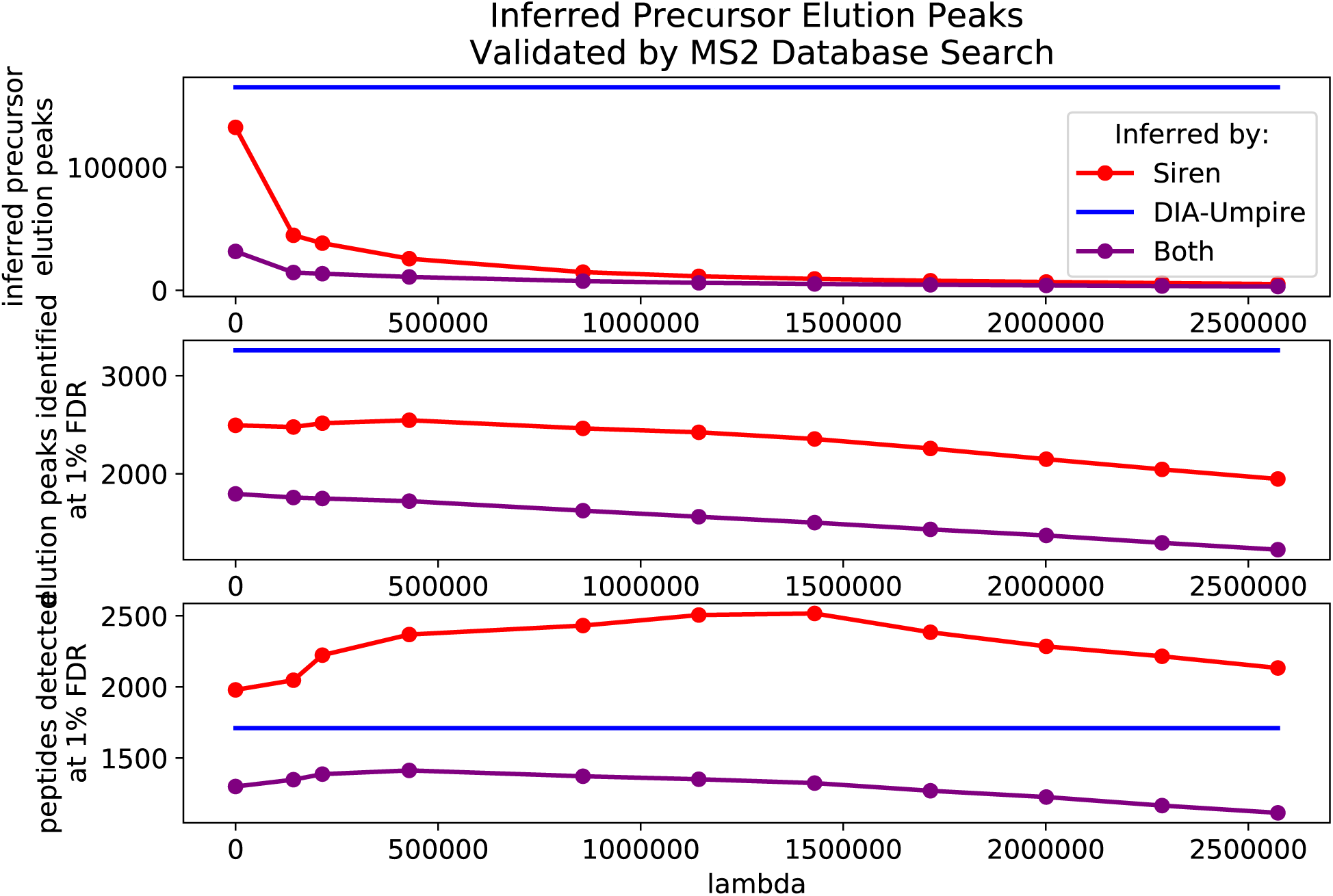
**Comparison of DIA-Umpire and Siren using raw MS2 spectra.** (A) Number of precursors annotated by DIA-Umpire and Siren, plotted as a function of Siren’s regularization parameter λ. The DIA-Umpire line is horizontal because it is invariant to λ. The purple line counts peaks that are identified by both methods, using a 0.01 m/z tolerance. (B) Number of PSMs accepted at 1% PSM-level FDR, using raw MS2 spectra with m/z values assigned by DIA-Umpire or Siren. (C) Number of peptides accepted at 1% peptide-level FDR.

The database search results show that the peaks produced by DIA-Umpire lead to fewer detected peptides than the peaks produced by Siren (Figure 4C). The unregularized (λ = 0) Siren model yields 1,979 detected peptides at 1% FDR, whereas DIA-Umpire detects 1,711 distinct peptides. Furthermore, tuning the regularization parameter λ yields further improvement for Siren, by preferentially eliminating low quality peaks and thereby reducing the multiple hypothesis testing burden during the database search step. A grid search of multiple values of λ led to peptide identifications as high as 2,516 peptides at 1% FDR (λ = 1429350) from 9,417 elution peaks.

Notably, the trend for the number of peptide-spectrum matches (PSMs) is reversed: at a 1% PSM-level FDR, DIA-Umpire accepts 3,529 PSMs, whereas Siren accepts at most 2546. This observation suggests that DIA-Umpire is assigning peaks to the same peptide at multiple points during its elution, or is identifying the same peptides in multiple charge states.

We also note that the low overlap between peaks produced by DIA-Umpire and Siren is maintained in the PSM-level and peptide-level results. This is perhaps not surprising, because as noted above, we expect the statistical power of detection to be low due to the use of raw MS2 DIA spectra in the database search step.

### 3.3 Comparison of Siren and DIA-Umpire using deconvolved MS2 spectra

To improve the MS2 analysis step of the comparison, we next compared Siren and DIA-Umpire in the context of DIA-Umpire’s full fragment deconvolution pipeline. To do this, we modified DIA-Umpire so that its fragment deconvolution step could accept as input elution profiles provided by Siren. In this way, we produced two comparable sets of pseudospectra, based on MS1 analysis by Siren and by DIA-Umpire. Subsequently, peptides were identified from the pseudospectra via database search. We carried out this analysis on both replicates of the Tsou2016 data set, as well as two additional data sets, derived from human HeLa cells [12] and from human spinal cord tissue [2]. For these analyses, we include decoy precursors in Siren (Section 2.1.5) to facilitate estimation of the FDP among the precursor peaks.

Across all of these data sets, and across a range of nominal precursor FDP values, we find that Siren detects more peptides than DIA-Umpire at an FDR threshold of 1% (Figure 5). At high FDP, Siren + DIA-Umpire result in roughly similar numbers of detected peptides; however, filtering some precursors using Siren’s λ parameter to achieve FDPs of 7.5-15% yields more identified peptides than DIA-Umpire’s full pipeline across all four data sets. An FDP of about 10% is close to optimal for each data set, suggesting that 10% may be a reasonable cutoff for more general use. At this threshold Siren improves upon DIA-Umpire by 7-26% across the four data sets. Notably, a significant proportion of the detected peptides are not shared between the two precursor annotation methods, suggesting that analysis of both sets of precursors jointly may be useful.

**Figure 5:**
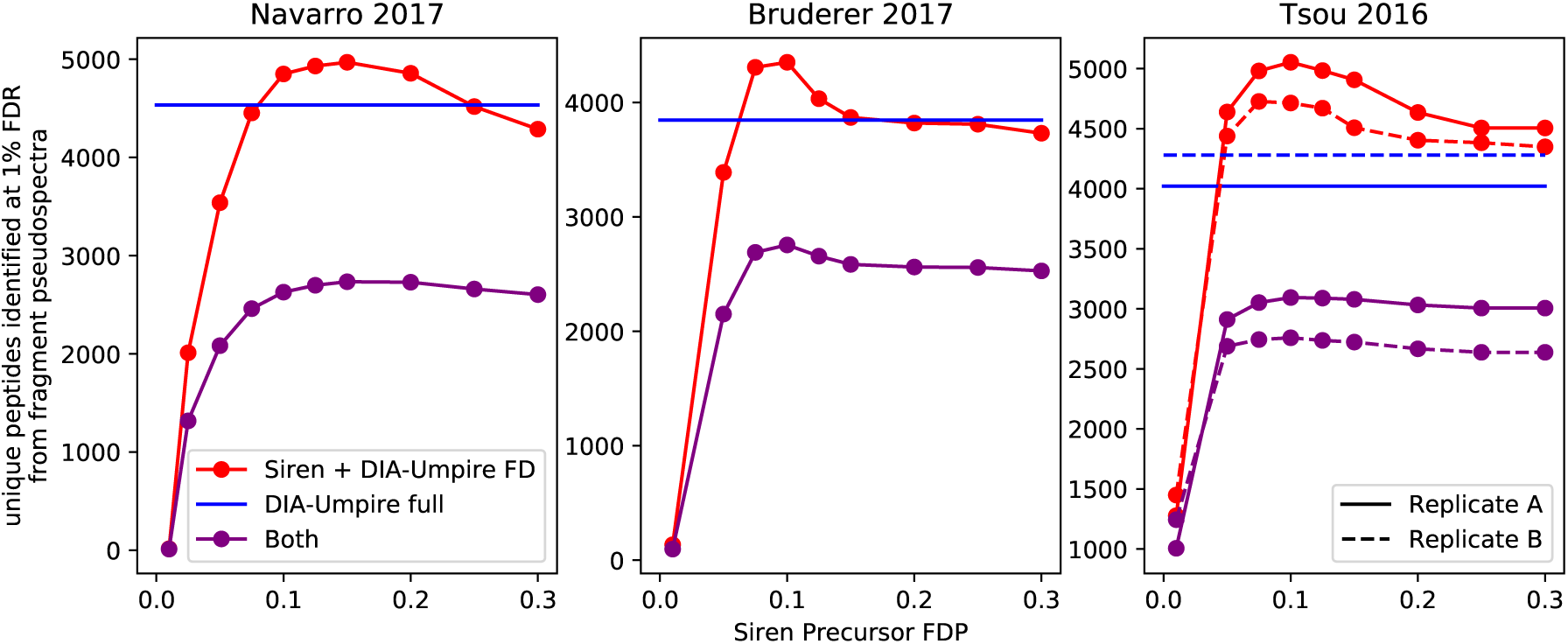
**Number of peptides identified by Siren and fragment deconvolution by DIA-Umpire.** The full pipeline of DIA-Umpire (DIA-Umpire full) and a pipeline consisting Siren and DIA-Umpire’s fragment deconvolution step (Siren + DIA-Umpire FD) were used to identify peptides via database search in four data sets. For Siren + DIA-Umpire FD, sets of peptide-spectrum matches were filtered by the false-discovery proportions (FDP) of their precursors, estimated by Siren. (0.01 m/z tolerance).

Because Siren’s regression model jointly considers the contributions of multiple precursors to observed peaks, the method is able to detect precursors even if their peaks exhibit interference. Figure 6 shows an example of this phenomenon. The two precursor ions (I and II) exhibit interference. Both Siren and DIA-Umpire confidently assign a peptide to precursor I, whose elution peak occurs around scan 613, but only Siren identifies precursor II, whose elution peak occurs around scan 617. In this case, Siren’s ability to detect the second precursor may be because precursor II’s second isotopic peak does not correlate well over time with the first and third due to interference from precursor I. Figure 6B shows that Siren was able to deconvolve the interference and recognize the elution profile of I.

**Figure 6:**
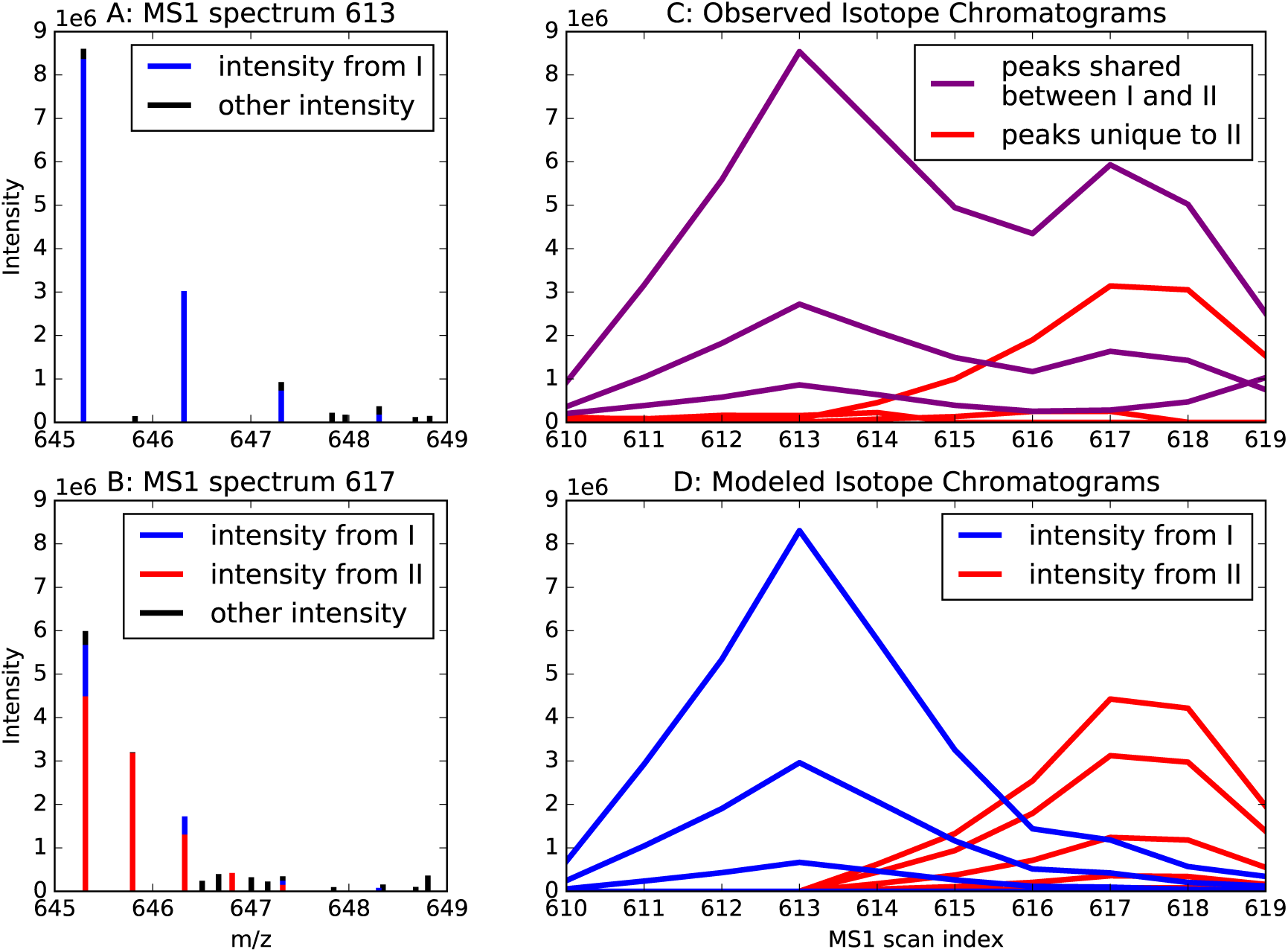
Peptide interference. (A) The figure plots a section of MS1 scan 613, depicting peaks identified by both DIA-Umpire and Siren as the isotope distribution of precursor ion *I* with monoisotopic m/z is 645.3 and charge +1. The colors denote the precursor ions contributing to peak intensity according to Siren. (B) A section of MS1 scan 617, depicting peaks identified Siren as the combined isotope distributions of precursor ion *I* and precursor ion *II*, both with monoisotopic m/z is 645.3 and and charges of +1 and +2, respectively. The colors denote the precursor ions contributing to peak intensity according to Siren. Siren suggests that the monoisotopic peaks of *I* and *II* interfere with each other, and that the second isotopic peak of *I* interferes with the third isotopic peak of *II.* Precursor *II* is not recognized by DIA-Umpire, but the corresponding MS2 spectrum is matched by database search to the sequence DISTNYYASQK. (C) Chromatograms of the ions of precursors *I* and *II.* Each chromatogram is labeled purple if its m/z value matches both precursors *I* and *II* or red if it matches only to precursor *II.* (D) Ion chromatograms modeled by Siren for precursors *I* and *II*, with colors corresponding to their precursors.

## 4. Discussion

Siren uses a regularized regression model to jointly infer precursor information at a scale suitable for DIA analysis. Although Siren’s model is similar to that of Nitpick, Siren’s sparse binning scheme allows it to scale to joint analysis of a full run. Like DIA-Umpire, siren infers precursor elution profiles that facilitate downstream peptide identification from MS2 spectra, but jointly models precursors such that it can deconvolve interference.

A key parameter to Siren is the regularization parameter λ. Siren employs a LASSO path estimation algorithm that simultaneously infers solutions for all values of λ. Subsequently, the user can choose a value of λ appropriate for the intended downstream use. Siren’s target-decoy strategy provides a mapping from λ values to estimated FDP, to assist in selecting λ. For instance, if the desired output is the accurate quantification of a peptide of known elution time, then a lower value of λ should be used. On the other hand, if the goal is to minimize false positives, then a higher value of λ may be more appropriate.

The inferred elution profiles in *B* can be used in many ways in both DDA and DIA analysis. The first benchmarking scheme (Section 3.2) suggests that Siren may be useful in selecting precursor isolation window in the context of DDA database search. The second benchmarking (Section 3.3) uses Siren’s inferred elution profiles to improve DIA analysis pipeline. Siren’s modeling of MS1 spectra could also be used in combination with a tool such as Specter [14], which models MS2 DIA spectra, so that both dimensions of data could be used simultaneously.

One challenge for Siren is that the model implicitly assumes that the observed data can be modeled as a mixture of peptide isotope distributions. Thus, non-peptide species in the sample will be modeled as noise in Equation 1. A direction for future work would be to incorporate into Siren a more sophisticated noise model that accurately captures various sources of noise in real MS1 data, thereby allowing Siren to focus its modeling on the signal induced by peptides.

## Acknowledgments

This work was funded by National Institutes of Health awards R01 GM121818 and P41 GM103533.

